# Multivariate white matter microstructure alterations in older adults with coronary artery disease

**DOI:** 10.1101/2025.04.25.650626

**Authors:** Stefanie A. Tremblay, Zacharie Potvin-Jutras, Dalia Sabra, Ali Rezaei, Safa Sanami, Christine Gagnon, Brittany Intzandt, Amélie Mainville-Berthiaume, Lindsay Wright, Ilana R. Leppert, Christine L. Tardif, Christopher J. Steele, Josep Iglesies-Grau, Anil Nigam, Louis Bherer, Claudine J. Gauthier

**Author notes:** Contributed equally to this work (Stefanie A. Tremblay, Zacharie Potvin-Jutras).

## Abstract

Patients with coronary artery disease (CAD) face an increased risk of cognitive impairment, dementia, and stroke. While white matter (WM) lesions are frequently reported in patients with CAD, the effects on WM microstructure alterations remain largely unknown. We aimed to identify WM microstructural alterations in individuals with CAD compared to healthy controls (HC), and to examine their relationships with cognitive performance. Forty-three (43) patients with CAD and 36 HC aged 50 and older underwent comprehensive neuropsychological testing and multi-modal 3T MRI. A novel multivariate approach - the Mahalanobis distance (D2) - was used to quantify WM abnormalities as the amount of deviation from the HC reference group. D2 integrates multiple MRI-derived diffusion-weighted imaging, R1 relaxometry, and magnetization transfer imaging metrics, while accounting for covariance between metrics. Relationships between WM D2 and cognition (executive function and processing speed) were also assessed. Compared to HCs, patients with CAD had higher D2 values in the whole WM (p=0.009) and in the territories right anterior, bilateral middle, and right posterior cerebral artery territories (p<0.05). Myelin-sensitive metrics, particularly R1 relaxation rate and MT saturation (MTsat), were the most important contributors to D2. Processing speed was positively associated with greater R1 in both the whole WM and left middle cerebral artery territory. These findings suggest that greater WM microstructural alterations observed in patients with CAD were mainly driven by differences in myelin content, as R1 and MTsat were the most important contributors. These alterations may contribute to a heightened risk of cognitive impairment.

## INTRODUCTION

Patients with coronary artery disease (CAD) are at an increased risk of cognitive decline, dementia, and stroke (Zheng et al., 2012; Kovacic et al., 2012; Olesen et al., 2017, 2024; Justin et al., 2013). This heightened vulnerability stems in part from cerebrovascular dysfunction, which affects cerebral vessels, grey matter and white matter (WM) (Barekatain et al., 2014; Launer et al., 2015; Vuorinen et al., 2014; Johansen et al., 2021). Major cardiac events, such as myocardial infarction, may further accelerate brain damage and the ensuing cognitive decline (Schievink et al., 2022; Xie et al., 2019). However, the underlying mechanisms linking CAD to cognitive decline remain poorly understood, hindering early detection and intervention efforts.

WM abnormalities, particularly WM lesions that appear as WM hyperintensities (WMHs) in magnetic resonance imaging (MRI), are frequently reported in CAD patients (Johansen et al., 2021; Vidal et al., 2010; Vuorinen et al., 2014). Because WM has lower perfusion than grey matter, it is especially vulnerable to changes in perfusion, and thus more prone to hypoxic injury as a result of cerebrovascular dysfunction (Inoue et al., 2023). These injuries are thought to follow vascular patterns, often occurring within specific arterial territories that are more susceptible to transient hypoperfusion. WMHs are known to contribute to the cognitive impairments seen in CAD patients (O’Brien, 2014; Filley & Fields, 2016). However, lesions and macrostructural measures of brain atrophy do not suffice to fully explain the impact of CAD on cognition as shown by Zheng and colleagues (2012), where CAD remained significantly associated with cognition after adjusting for WMH, and hippocampal and cortical GM volumes (Santiago et al., 2015). This suggests that more sensitive neuroimaging metrics are needed to detect subtle changes in brain health not captured by lesions and volumetry, such as microstructural changes in the so-called “normal-appearing” WM. Diffusion-weighted imaging (DWI), magnetization transfer imaging (MTI) and relaxometry allow to capture such changes in WM microstructure.

Changes in diffusion tensor (DTI) metrics in several major WM tracts, including the fornix, body of the corpus callosum, superior corona radiata and superior fronto-occipital fasciculus, have been reported in CAD patients (Poirier et al., 2024). Importantly, WM microstructural integrity, quantified using fractional anisotropy (FA), has been linked with cognitive performance, especially executive function and processing speed, in cognitively intact CAD patients (Santiago et al., 2015). This suggests that alterations in WM microstructure may be an important factor contributing to subtle changes in cognition in CAD patients. However, the mechanisms through which WM damage occurs and causes cognitive decline are poorly understood, as the few studies that have looked at this used techniques that are physiologically unspecific (i.e., DTI) (Poirier et al., 2024; Santiago et al., 2015; Riffert et al., 2014). Metrics derived from more advanced multi-compartment DWI models to assess volumes of axonal and extra-cellular components, complemented with quantitative measures of tissue composition including myelin density (MTsat and R1), can provide mechanistic insights into the pathophysiological mechanisms underlying WM microstructural alterations in CAD patients.

In this cross-sectional study, we aimed to identify WM microstructural abnormalities in patients with CAD and to examine their relevance to cognitive function. We employed a multivariate approach using the Mahalanobis distance (D2), which quantifies the extent of deviation between an individual and a reference distribution (e.g., healthy controls) across multiple MRI measures (Tremblay et al., 2024), to compare WM microstructure between CAD patients and healthy controls. This approach yields a voxel-wise index of abnormality that integrates several microstructural metrics while accounting for their covariance. We examined deviations in WM microstructure within atlas-based arterial territories (Liu et al., 2023), as WM damage in vascular disease often results from transient hypoperfusion and hypoxia—processes that typically affect specific vascular regions (Inoue et al., 2023; O’Rourke & Hashimoto, 2007). Finally, we assessed whether these WM abnormalities were associated with cognitive performance, focusing on executive function and processing speed—domains frequently impaired in CAD.

## METHODS

### Participants

Ninety-nine (99) participants of 50 years and above were recruited, of which 87 completed the study (46 CAD patients and 41 healthy controls; HC). Out of 12 participants that dropped out, five participants did not complete the study due to interruptions during the COVID-19 pandemic, two participants were excluded due to claustrophobia, two participants were excluded due to discomfort during the MRI and three participants dropped out for personal reasons (e.g. loss of interest, issues with scheduling). Because the MRI acquisition also included a hypercapnia manipulation (not described or used here), which can cause discomfort in some individuals, our attrition rate was higher than that of conventional MRI sessions. The study was approved by the Ethic review board of the Montreal Heart Institute, in accordance with the Declaration of Helsinki. Written informed consent was obtained at the first visit. A medical questionnaire, which had been previously filled out on the phone, was also reviewed at this visit and the mini-mental state examination (MMSE) was administered to ensure eligibility (see exclusion criteria).

Inclusion criteria for patients included documented coronary artery disease (prior acute coronary syndrome, prior coronary angiography or revascularization, or myocardial ischemia documented on myocardial scintigraphy). Healthy controls (HCs) had to be free of any cardiac and neurological issues, diabetes and hypertension. All participants had to be fluent in either English or French (for informed consent and cognitive assessment).

Exclusion criteria for all participants included history of stroke, neurological, psychiatric or respiratory disorders, thyroid disease, potential cognitive impairment (MMSE < 25), tobacco use, high alcohol consumption (more than 2 drinks per day), contraindications to MRI (e.g., ferromagnetic implant, claustrophobia), and use of oral or patch hormone therapy. Participants were also excluded if they had surgery under general anesthesia within the last 6 months, a recent acute coronary event (< 3 months), chronic systolic heart failure, resting left ventricular ejection fraction < 40%, symptomatic aortic stenosis, severe nonrevascularizable coronary artery disease, including left main coronary stenosis, awaiting coronary artery bypass surgery, implanted automatic defibrillator or permanent pacemaker. Excessive discomfort due to hypercapnia (> 5 on the dyspnea scale of Banzett et al., 1996) also constituted an exclusion criterion.

All participants completed a neuropsychological battery and an MRI session. Participants who had all DWI and MTI data were included in this study (N= 84). However, 5 of these participants were excluded due to the presence of artefacts in their DWI data (N=4) or due to an incidental finding in their MRI data (N=1), resulting in a final sample size of 79. Of those, 43 were CAD patients (age = 68.2 ± 8.7 years, 8 females) and 36 were HCs (age = 64.1 ± 7.8, 10 females).

Detailed clinical characteristics of CAD patients can be found in Table 1. Briefly, the CAD group consisted of patients diagnosed with coronary artery disease. Almost half of the patients (18) had a history of myocardial infarction, 65% underwent a percutaneous coronary intervention (i.e., stent) and 23% had a coronary artery bypass grafting. CAD patients were treated with lipid-lowering medications, beta blockers, aspirin, renin-angiotensin-aldosterone system (RAAS) inhibitors, and calcium channel blockers.

**Table 1.** Clinical Health Characteristics of patients with CAD (n=43) and HCs (n=36)

| Groups | Patients with CHD (n=43) | Healthy Controls (n=36) |
| --- | --- | --- |
| **Cardiometabolic risk factors, n (%)** |  |  |
| Type 2 Diabetes | 6 (7) | 0 |
| Hypertension | 23 (53) | 0 |
| Dyslipidemia | 35 (81) | 1 (3) |
| Obesity | 11 (26) | 5 (14) |
| **Medication, n (%)** |  |  |
| Beta-Blocker | 21 (49) | 0 |
| Aspirin | 14 (33) | 0 |
| Antilipemic agents (e.g., statins) | 29 (67) | 1 (3) |
| RAAS inhibitor | 11 (26) | 0 |
| Calcium channel blockers | 7 (16) | 0 |
| **Cardiovascular history, n (%)** |  |  |
| Previous MI | 18 (42) | - |
| STEMI | 5 (12) | - |
| NSTEMI | 13 (30) | - |
| Percutaneous coronary intervention | 28 (65) | - |
| Coronary artery bypass grafting | 10 (23) | - |
| Angina | 13 (30) | - |
| Familial history of cardiovascular disease | 27 (63) | - |

*MI = Myocardial Infarction; STEMI = ST-elevation Myocardial Infarction; NSTEMI = Non-ST-elevation Myocardial Infarction; RAAS = Renin-Angiotensin-Aldosterone System*

### MRI Protocol

MRI data were acquired on a 3T Siemens Magnetom Skyra scanner at the Montreal Heart Institute. The multi-shell DWI acquisition was a pulse gradient spin-echo sequence with echo planar imaging (EPI) readout (TR= 6000 ms, TE= 106 ms, phase-encoding direction = posterior-anterior (PA), resolution = 2 mm isotropic) across 3 diffusion-weighted shells with gradient strengths of b= 300 (10 directions), 700 (30 directions), and 2500 s/mm^2^ (64 directions), and 3 volumes acquired without diffusion weighting (b = 0). Six non-diffusion weighted volumes (b = 0) were also acquired in the opposite phase encoding direction (AP) for distortion correction.

Three gradient echo sequences (TR= 33 ms, TE= 3.81 ms, flip angle= 10°, resolution= 2 mm isotropic), one with (MT-w) and one without a preparatory MT pulse (MT-off), and a T1w image (TR= 15 ms, TE= 3.81 ms, flip angle= 25°, resolution= 2 mm isotropic) were acquired for MTsat computation. An off-resonance MT pulse (off-resonance frequency = 2.2 kHz, duration = 12.8 ms, flip angle = 540°) was applied prior to RF excitation to obtain MT-weighting (Helms et al., 2008). A B1 map (an anatomical image and a flip angle map) (TR = 19780 ms, TE= 2.36 ms, flip angle = 8°) was also acquired using a TurboFLASH with a preconditioning RF pulse to correct B1 field inhomogeneities (Chung et al., 2010). This custom pulse sequence was provided by the McConnell Brain Imaging Centre of the Montréal Neurological Institute.

High-resolution T1-weighted structural images were acquired with a Magnetization Prepared RApid Gradient Echo (MPRAGE) sequence (TR= 2300 ms, TE= 2.32 ms, flip angle= 8°, resolution= 0.9 mm isotropic). These images were used for segmentation by tissue type.

### Preprocessing

We derived 12 microstructural metrics from the DWI and MTI data. These metrics were obtained from the diffusion tensor imaging (DTI) model, the fixel-based analysis (FBA) framework—which estimates fibre density and cross-section from fibre orientation distributions (FODs) computed using multi-tissue constrained spherical deconvolution (CSD) (Jeurissen et al., 2014), and the neurite orientation dispersion and density imaging (NODDI) model (Zhang et al., 2012). The MT saturation was computed from the MT-w and MT-off images, using the T1w images to reduce T1 dependence (Helms et al., 2008), and R1 (1/T1) maps were also generated.

#### Diffusion Tensor Imaging

Most preprocessing was performed using the MRtrix3 toolbox (v3.0.2) (Tournier et al., 2019). DWI data underwent denoising (`dwidenoise`) and correction for motion, eddy currents, and susceptibility-induced distortions using `dwifslpreproc`, which leverages the Eddy and `topup` tools in FSL (v6.0.1) (Andersson et al., 2003; Andersson & Sotiropoulos, 2016; Smith et al., 2004). `Topup` corrected susceptibility-induced distortions by utilizing pairs of b0 volumes with opposite phase-encoding polarities (AP) and with the same phase encoding as the input DWI series. The preprocessed DWI data were then upsampled to match the MPRAGE T1w image resolution (0.9 mm isotropic). A brain mask was generated using FSL’s brain extraction tool (`bet`) and applied to eliminate non-brain voxels from the DWI images (Jenkinson, 2005). Bias field correction was performed using the N4 algorithm in ANTs (v3.0) (Tustison et al., 2010), after which the tensor was computed on the corrected DWI data using `dwi2tensor`. DTI metrics—including fractional anisotropy (FA), mean diffusivity (MD), axial diffusivity (AD), and radial diffusivity (RD)—were then extracted using `tensor2metric` (Basser et al., 1994).

#### Fixel-based analysis

The FBA pipeline was used to derive fibre density and cross-section from FODs (Tournier et al., 2019). MPRAGE T1-w images were rigidly and linearly registered to the non-diffusion weighted (b=0) preprocessed DWI volume using `antsRegistration` (v2.4.2). These images were subsequently segmented using the `5ttgen` FSL function in Mrtrix3, based on the FAST algorithm (Avants et al., 2009; Smith et al., 2012). Response functions for white matter (WM), grey matter (GM), and cerebrospinal fluid (CSF) were then computed from the preprocessed DWI data (without N4 bias field correction) and the five-tissue-type (5tt) image using `dwi2response` (`msmt_5tt` algorithm) (Jeurissen et al., 2014). Bias field correction was deferred to a later stage of the FBA pipeline (Raffelt et al., 2017). The computed WM, GM, and CSF response functions were averaged across all participants to obtain a single response function for each tissue type. Multi-shell multi-tissue CSD was performed using the averaged response functions to estimate orientation distribution functions (ODFs) for each tissue type (Jeurissen et al., 2014), implemented with the `dwi2fod msmt_csd` function within the brain mask. Bias field correction and global intensity normalization were then applied to the ODFs using the `mtnormalise` function to ensure comparability across subjects function (Raffelt et al., 2017).

#### Registration

To optimize WM and GM alignment, multi-contrast registration was performed. Population templates were created from WM, GM, and CSF FODs and brain masks of all participants using the `population_template` function in Mrtrix3 (with regularization parameters: nl_update_smooth=1.0 and nl_disp_smooth=0.75), generating a group template for each tissue type (Tournier et al., 2019).

Subject-to-template warps were computed using `mrregister` with the same regularization parameters and applied to brain masks, WM FODs, and DTI metrics (i.e., FA, MD, AD and RD) using `mrtransform` (Raffelt et al., 2011). At this stage, WM FODs were transformed but not reoriented; the voxels of the images were thus aligned but not the fixels (“fibre bundle elements”). A template mask was created as the intersection of all warped brain masks (`mrmath min` function), including only voxels present in all subjects. The WM volumes of the five-tissue-type (5tt) 4D images were also warped to the group template space and averaged across participants to generate a WM mask for analyses.

#### Computing fixel metrics

The WM FOD template was segmented into fixels using the `fod2fixel` function (Raffelt et al., 2011; Smith et al., 2013), defining the fiber bundle elements to be analyzed within each voxel. Fixel segmentation was performed on each subject’s WM FODs using `fod2fixel`, yielding the apparent fibre density (FD) metric.

The `fixelreorient` and `fixelcorrespondence` functions ensured accurate mapping of subjects’ fixels onto the fixel mask (Tournier et al., 2019).

Fibre bundle cross-section (FC) was computed from the warps generated during registration (using `warp2metric`), quantifying the expansion/contraction needed to align individual fixels with the template. Lastly, fibre density and cross-section (FDC), representing the total information-carrying capacity of a fibre bundle, was computed as the product of FD and FC.

#### Transforming fixel metrics into voxel space

To integrate all metrics into a multi-modal model, fixel metrics were transformed into voxel-wise maps. For fiber density, we used the *l* = 0 term of the WM FOD spherical harmonic expansion (i.e., the first volume of the WM FOD, equal to the sum of FOD lobe integrals), yielding a total fibre density (FD total) per voxel. This method provides more reproducible estimates than summing the FD fixel metric (Calamante et al., 2015). The FOD *l* = 0 term was scaled by the spherical harmonic basis factor (multiplying voxel intensities by the square root of 4π).

Fiber cross-section was aggregated by computing the mean FC weighed by FD (using the `mean` option of the `fixel2voxel` function), capturing the typical expansion/contraction required to align fiber bundles within a voxel. Finally, the voxel-wise sum of FDC, representing total information-carrying capacity per voxel, was computed using `fixel2voxel` with the `sum` option.

#### NODDI metrics

Bias field-corrected DWI data were fitted to the NODDI model using the Python implementation of Accelerated Microstructure Imaging via Convex Optimization (AMICO) (Zhang et al., 2012; Daducci et al., 2015). A diffusion gradient scheme file was generated from the bvecs and bvals files. Response functions were computed for all compartments, and model fitting was performed within the brain mask. The extracted parameters included the intracellular volume fraction (ICVF, also called neurite density), the isotropic volume fraction (ISOVF), and the orientation dispersion index (OD). These NODDI metrics were subsequently warped to group space using previously computed transforms.

#### MTsat and R1 maps

The MTsat and R1 maps were calculated using the hMRI-toolbox (v0.3.0) in Statistical Parametric Mapping (Tabelow et al., 2019). The anterior and posterior commissure of the MT-w, MT-off and T1w images were reoriented to MNI space. The reorientation process was performed before processing the MTsat and R1 maps to enhance the alignment of the images’ WM. The B1 transmit bias correction field was computed using an anatomical map and a scaled flip angle map from a turbo flash sequence using the “Create hMRI maps” module of the hMRI-toolbox. Then, the B1 correction field, MT-w, MT-off and T1w images were included in the “Create hMRI maps” module to measure MTsat and R1 maps with default parameters. Finally, non-brain voxels were removed from the T1w maps using FSL’s brain extraction tool (bet). The T1w brain masks were then applied to the MTsat and R1 maps. The MTsat and R1 maps were registered and warped (rigid, linear) using the MPRAGE image that was previously warped to the DWI space (within-subject) as a target, with `antsRegistration`. The MTsat and R1 maps were then warped to the population template using the previously generated transforms.

### Computing multivariate distance metric (D2)

D2 was computed from the 12 WM features (FA, AD, RD, MD, ICVF, ISOVF, OD, AFDtotal, meanFC, sumFDC, MTsat and R1) using the MVComp toolbox (Tremblay et al., 2024). Prior to D2 calculation, the effect of age was removed from feature maps by fitting a linear model predicting voxel-wise metric values from age using `LinearRegression` in `sklearn.linear_model` and computing the residuals. Residualized maps were then used for D2 calculation.

D2 was computed voxel-wise between each subject of the CAD group and the reference, consisting of the group average of the HCs. The D2 scores obtained thus indicate the extent of deviation in WM microstructure from a healthy reference. Group averages for each WM metric were computed from the reference group (N = 36) using MVComp’s `compute_average` function. The covariance matrix (s) and its pseudoinverse (pinv_s) were derived from the reference data using the `norm_covar_inv` function. To visualize the relationships between MRI metrics, a correlation matrix was generated using the `correlation_fig` function, which calculates correlations based on the covariance matrix (s) (Fig 1). D2 was then computed within MVComp using the following equation:

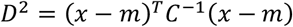

where *x* represents the data vector for an individual observation (e.g., one subject), *m* is the vector of group-averaged values for each MRI metric, and *C*^*-1*^ is the inverse of the covariance matrix. The `model_comp` function was used to compute voxel-wise D2 between each subject and the reference average within a specified analysis mask. Here, a WM mask generated from the averaged WM volumes of the five-tissue-type images across all participants was provided. To minimize partial volume effects, the threshold was set at 0.97. The `model_comp` function yields a matrix containing D2 values for all subjects, with dimensions corresponding to the number of voxels x number of subjects. Finally, the `dist_plot` function was used to generate individual D2 maps in nifti format. The overall workflow for D2 computation is illustrated in Figure 1.

**Figure 1.**
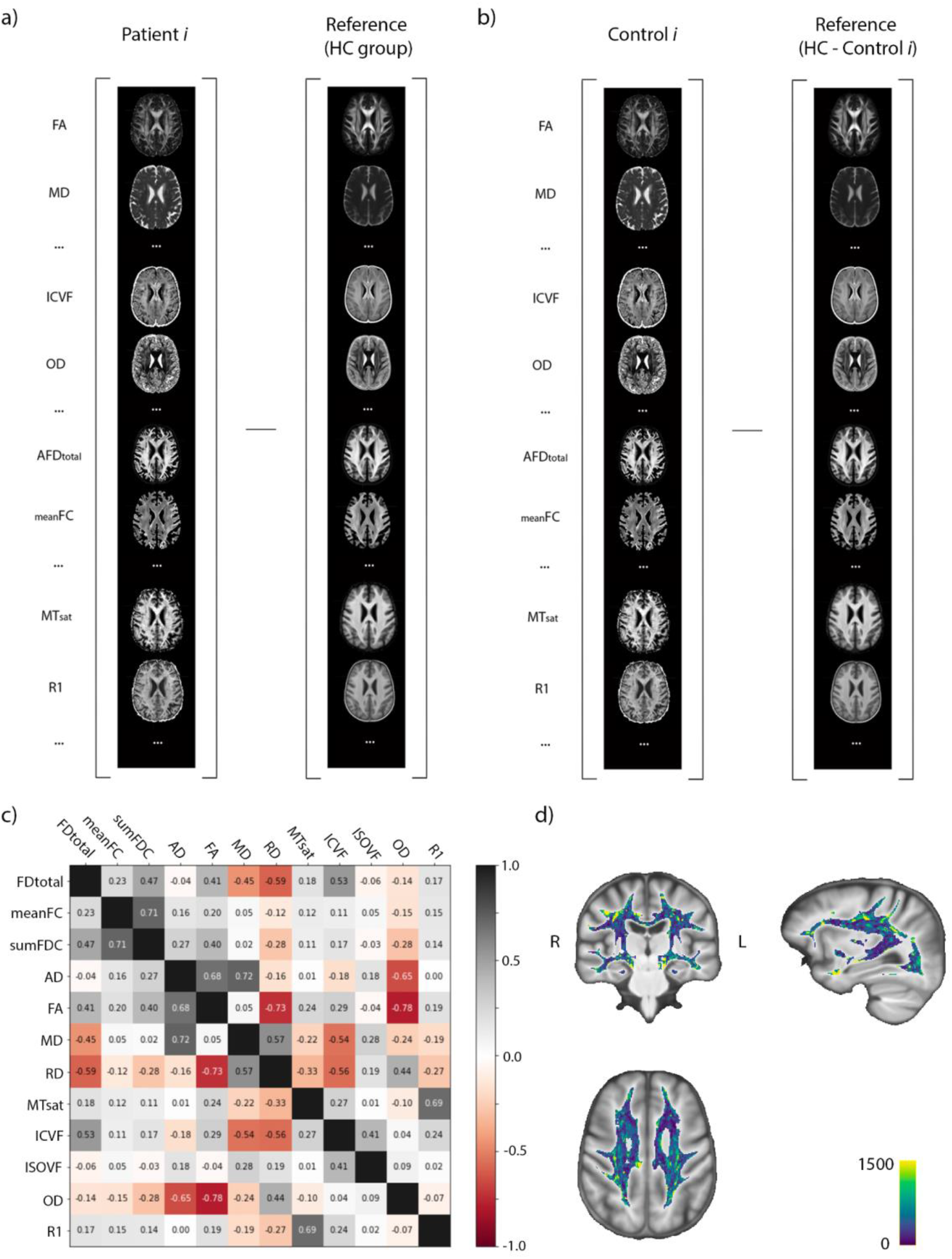
Workflow for computing D2. a) For subjects of the CAD group, D2 is computed by subtracting the data of each patient i and the reference sample, here the average of the HC group (N= 34). b) For subjects of the HC group, D2 is computed by subtracting the data of each control i and the reference sample, here the average of the HC group excluding the subject under evaluation (HC group - control i). c) The correlation matrix shows relationships between MRI metrics (residualized for age), highlighting the importance of accounting for covariance between variables in multivariate frameworks. d) Example D2 map of a patient. The intensity indicates the amount of deviation in the WM microstructure of this subject relative to the reference, at each voxel.

D2 maps were also computed for subjects of the HC group using a leave-one-out approach to exclude the subject under evaluation (i.e., comparing each subject of the HC group to a personalized reference group that includes all other HCs but excludes the subject under study). This is done within MVComp using the `exclude_comp_from_mean_cov` option of the `model_comp` function.

Because we wanted the reference to represent a healthy state, subjects of the HC group that were identified as outliers in D2 (i.e., mean D2 in whole WM > 2 SD) were excluded from the reference and from further analyses (N= 2). D2 was re-computed with this new reference (N= 34) for both the CAD and HC groups (using the leave-one-out approach).

### Extracting D2 in arterial territories

A cerebral arterial territories atlas (Liu et al., 2023) was used to define regions of interests (ROIs) in WM (**Supplementary Figure 1**). The 3D MR-based cerebral arterial atlas (Liu et al., 2023) was registered and warped to the D2 maps space using `antsRegistration`. Average D2 values were then calculated across WM voxels in the left and right anterior cerebral artery (ACA) territory, middle cerebral artery (MCA) territory and posterior cerebral artery (PCA) territory.

### Determining features contribution to D2

The relative contributions of each feature (i.e., MRI metric) to D2 in the whole WM, as well as in each arterial territory, were extracted using the `return_raw` option of the `model_comp` function in MVComp. The return_raw option yields a matrix of size (number of voxels) x (number of metrics) x (number of subjects). Contributions were then summarised by averaging distance values across voxels within the mask and across subjects and dividing by the total distance (for all features), resulting in one distance value per metric, expressed as a percentage, for each region. This analysis provides a measure of the importance of each metric in determining D2.

### Neuropsychological assessment

A comprehensive neuropsychological assessment was administered in the following order. The Montreal Cognitive Assessment (MoCA) was used to assess global cognition, the Digit Symbol Substitution Test (DSST) as a measure of processing speed. Then, the Delis-Kaplan Executive Function System (D-KEFS) Color Word Interference Test (CWIT) test, which consists of 4 conditions (i.e., color naming, reading, inhibition and switching) that assess processing speed and different aspects of executive function, was conducted (Delis, Kaplan, & Kramer, 2001; Lezak, 2004). The Trail Making Test (TMT) part A, which measures processing speed, and part B, a measure of executive function, was administered last.

All scores were transformed into standardized z-scores and composite scores, representing different cognitive domains, were then created using those z-scores (Desjardins-Crepeau et al., 2014): (1) executive functioning = ((TMT B + CWIT 3 + CWIT 4 z-scores)/3)*-1; (2) processing speed = ((DSST*-1 + TMT A + CWIT 1 + CWIT 2 z-scores)/4)*-1. Chronbach’s alphas (α) were used to assess the reliability of the cognitive composite scores (see in Results) (Desjardins-Crepeau et al., 2014). Positive z-scores indicated more items completed in a given time and thus better performance in these cognitive domains.

### Statistical analyses

Group differences in demographic, hemodynamic measures, and cognitive measures between patients with CAD and HCs were tested using ANCOVA, controlling for age and sex (and for education in the case of analyses related to cognition). Binomial logistic regression was used to assess group differences in the frequency of males vs females. D2 maps were created from age-residualized feature maps, and should thus be free of any age effects. Age was also regressed out from composite scores of cognition (i.e., executive function and processing speed) and the residuals were used in all subsequent analyses.

To assess whether the amount of WM alteration differed in CAD compared to HCs, we performed ANCOVA analyses in each arterial territory and in the whole WM, adjusting for sex. We then extracted the contributions (i.e., loadings) of each WM metric to D2 within each region as described above. This allows the recovery of biological specificity, which facilitates the interpretation of findings (Tremblay et al., 2024, 2025).

Linear regression models were then used to assess the relationships between cognition (i.e., executive function and processing speed) and D2 in regions where significant group differences were found, with sex and education as covariates. Regression analyses were also performed between cognition and the WM metrics that contributed most to D2 (i.e., those that contributed ≥10%) as the direction of relationships with individual metrics is more readily interpretable biologically. A subject of the CAD group who was identified as an outlier (whole WM D2 > 2 SD from the mean) was excluded from those analyses.

We assessed whether the association between WM microstructure and cognition differed across groups by adding a group interaction term to significant models.

To account for multiple comparisons across tests and regions, p-values were adjusted using the Benjamini-Hochberg method, with a significance threshold set at 0.05 (equivalent to a p-value uncorrected of 0.008).

## RESULTS

Clinical health characteristics of patients with CAD can be found in **Table 1**.

### Group differences

There were significant group differences in several parameters, whereby patients with CAD exhibited higher BMI (p < 0.001; n^2^p = 0.139), waist circumference (p < 0.001; n^2^p = 0.185) and body fat mass (p < 0.001; n^2^p = 0.220) compared to HCs (Table 2). Other demographic parameters such as age, sex, education, hemodynamic measures and cognition did not differ between groups. The Chronbach’s alpha values indicated acceptable internal consistency, with α = 0.95 for processing speed and α = 0.82 for executive function.

**Table 2.** Demographic data, hemodynamic measures, and cognitive scores in both groups.

| **Demographic** | **Patients with CAD**<br>**(n=43)** | **Healthy Controls**<br>**(n=36)** | **p-values** |
| --- | --- | --- | --- |
| Age (years) | 68.2 ± 8.7 | 64.1 ± 7.8 | 0.180 |
| Sex (M/F) | 35 / 8 | 26 / 10 | 0.285 |
| Education (years) | 16.2 ± 4.5 | 16.0 ± 2.4 | 0.632 |
| BMI (kg/m<sup>2</sup>)* | **28.1 ± 4.5** | **26.0 ± 3.9** | **< 0.001** |
| Waist Circumference (cm)* | **103.0 ± 9.6** | **94.5 ± 11.8** | **< 0.001** |
| Body Fat (%)* | **29.2 ± 7.7** | **25.7 ± 7.2** | **< 0.001** |
| **Hemodynamic Measures** |  |  |  |
| Systolic BP (mmHg) | 126.9 ± 14.3 | 128.3 ± 14.0 | 0.732 |
| Diastolic BP (mmHg) | 73.3 ± 7.2 | 76.0 ± 8.8 | 0.653 |
| Resting Heart Rate (bpm) | 63.4 ± 10.9 | 68.6 ± 10.1 | 0.101 |
| **Cognition** |  |  |  |
| MoCA score | 25.4 ± 3.0 | 26.6 ± 2.2 | 0.061 |
| MMSE | 28.4 ± 1.3 | 28.2 ± 1.5 | 0.796 |
| Executive function (z-score) | - 0.24 ± 0.95 | 0.29 ± 0.63 | 0.066 |
| Processing speed (z-score) | - 0.21 ± 0.82 | 0.25 ± 0.70 | 0.068 |

*BMI = Body Mass Index; BP = Blood Pressure; MoCA = Montreal Cognitive Assessment; MMSE = Mini-mental State Examination*

Patients with CAD revealed significantly higher WM deviation from a healthy reference (D2) compared to HCs in the whole WM (p=0.009; n^2^p = 0.091), right ACA WM territory (p=0.005; n^2^p = 0.103), left MCA WM territory (p=0.047; n^2^p = 0.054), right MCA WM territory (p=0.024; n^2^p = 0.069) and right PCA WM territory (p=0.047; n^2^p = 0.054) (**Figure 2**).

**Figure 2.**
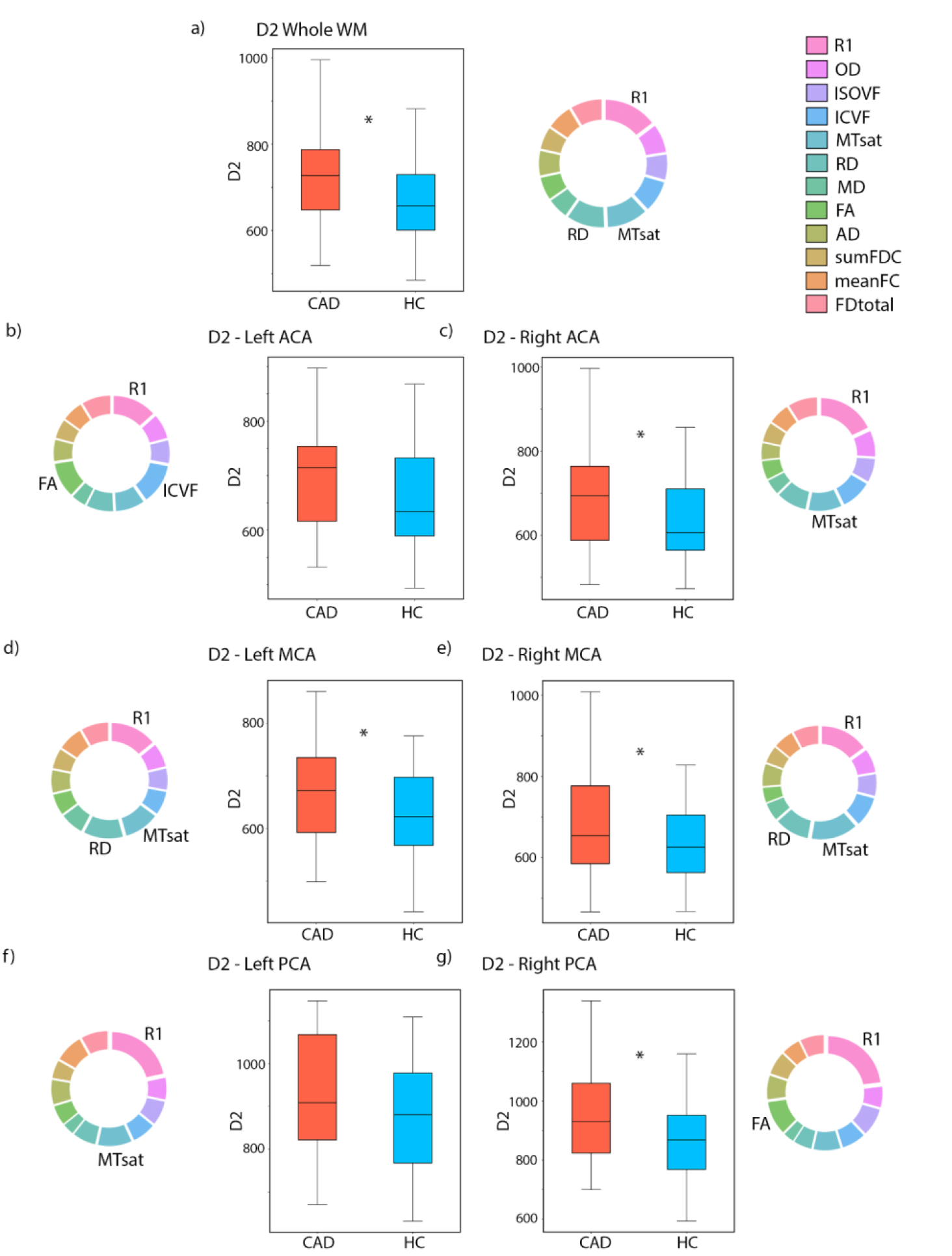
Box plots showing D2 values for each group (CAD in red; HC in blue) in the whole WM (a) and in each arterial territory (b-g). Significant group differences are indicated by an asterisk. Pie charts show the relative contribution (%) of each MRI metric to D2 in each region. The metric name is indicated only for the most important contributors (those that account for >10%), for clarity.

### Features’ contribution to D2

Overall, R1 was the most important contributor to D2 in all regions, ranging from contributions of 13.4% in the left ACA to 23% in the right PCA. Additionally, MTsat emerged as another important metric in four arterial territories and in the whole WM **(Figure 2)**. Percentage contributions of the top contributors in each region of interest can be found in **Figure 2** and are presented in **Supplementary Table 1**. Subsequent regression analyses with cognition focused on regions where significant group differences were found and on WM metrics with the highest loadings (>10%) within each region.

### Links between D2, WM metrics and cognition

There were no significant relationships between D2 and cognition (i.e., executive function and processing speed) in any region.

Relationships between cognitive performance and individual WM metrics were also assessed, focusing on metrics that contributed significantly (>10%) to D2 within each region. Processing speed was positively associated with R1 in the whole WM (R^2^ = 0.213; p = 0.008), in the right ACA (R^2^ = 0.181 ; p = 0.038), and in the left MCA WM territory (R^2^ = 0.235; p = 0.003) in the whole sample. Processing speed scores were also positively associated with MTsat in the right ACA territory (R^2^ = 0.184 ; p = 0.032). Only the correlations between processing speed and R1 in the whole WM and R1 in the left MCA remained significant after FDR correction (*p*_FDR_ < 0.05 or *p* uncorrected ≤ 0.008) (**Figure 3**). In both of these regression analyses, the covariates sex and education were found to be significant (p < 0.05). However, no significant group interactions were observed (p = 0.063 in the whole WM and p = 0.073 in the left MCA WM territory).

**Figure 3.**
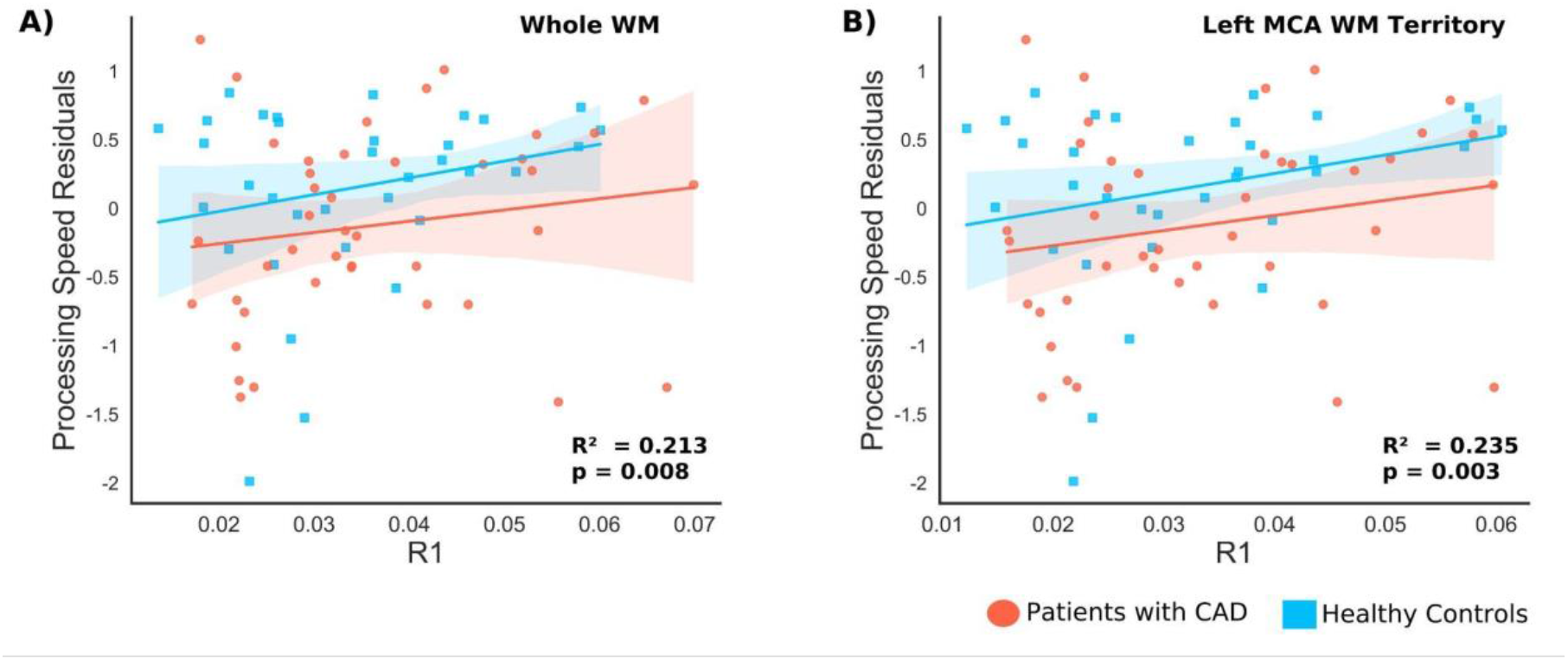
Associations between R1 and processing speed residuals (z-scores) in A) the whole WM and B) the left middle cerebral artery (MCA) WM territory in the whole sample.

There were no significant associations between individual WM metrics and executive function (*p* > 0.05).

## DISCUSSION

In this study, we documented the presence of WM microstructural alterations in CAD patients as compared to HCs, and a link between R1, the most important driver of microstructural alterations, and cognition. Deviations in WM microstructure were quantified using a novel multivariate approach that allows the integration of several MRI measures into a single score indicative of overall WM abnormality. Patients with CAD displayed significantly higher D2 values in the right ACA territory, bilateral MCA territory, right PCA territory, and whole WM compared to HCs. Additionally, myelin content emerged as a primary mechanism underlying these alterations, with R1 and MTsat identified as important contributors to D2 in most regions. Processing speed was positively associated with R1 in the left MCA territory and whole WM in the whole sample, but there were no significant group differences in these effects. In summary, these findings suggest that patients with CAD exhibit significant and complex WM microstructure alterations compared to HCs, with myelination being most affected. Furthermore, our data indicates a relationship between higher processing speed and greater macromolecular content.

### WM microstructural alterations in CAD patients

Cardiovascular disease is known to impact several aspects of brain health, including WM (Barekatain et al., 2014; Launer et al., 2015; Vuorinen et al., 2014; Haight et al., 2018). However, most studies investigating WM changes in patients with CAD have focused on macrostructural changes in WM, such as the volume of WM lesions (WMH) (Johansen et al., 2021; Vidal et al., 2010; Vuorinen et al., 2014), while only a few studies have investigated WM microstructural health (Poirier et al., 2024; Santiago et al., 2015; Li et al., 2024). Here, we found that the WM microstructure of patients with CAD is significantly different from that of HCs in several arterial territories (right ACA, bilateral MCA, right PCA), as well as in the whole WM (Figure 2). Our findings revealed that R1 was the most important contributor to D2. This suggests that variations in myelin content, without concomitant variations in axonal content, were the main factor underlying WM alterations in CAD patients (Callaghan et al., 2014, Draganski et al., 2011, Helms et al., 2008). However, it is also worth noting that many other WM features contributed to D2, to varying degrees in different regions, highlighting complex and regionally-specific effects.

MTsat, RD, and FA were also identified as important contributors (>10%) in several regions. The strong contributions of MTsat and RD further support differences in myelin as a primary biological mechanism underlying WM alterations in individuals with CAD, particularly in the left ACA and bilateral MCA (Helms et al., 2008; Winklewski et al., 2018). In contrast, FA, a metric that can reflect several WM properties but that is most sensitive to changes in orientation dispersion, was a strong contributor to D2 in the left PCA (Zhang et al., 2012).

Our results build on previous studies on WM microstructure alterations in individuals with CAD (Poirier et al., 2024; Li et al., 2024). In the study by Poirier and colleagues (2024), patients with CAD exhibited differences in FA and MD, metrics that can reflect both fiber density and orientation dispersion but that are inherently non-specific, in WM tracts primarily perfused by the ACA and MCA. Similarly, a study using diffusion kurtosis imaging found reductions in tissue complexity and integrity across widespread WM regions (Li et al., 2024). Consistent with these findings, our results revealed WM microstructure disruption across all cerebral arterial territories in CAD. However, the strong contribution of R1 and MTsat to D2 suggest that myelin loss is likely a primary mechanism of WM pathological alteration in CAD. To our knowledge, no other studies have examined changes in myelination specifically in patients with CAD, although lower myelin content has been reported in older adults with cardiovascular risk factors such as hypertension and obesity (Trofimova et al., 2023).

Together, these findings highlight the importance of using a multivariate approach to assess WM health in CAD, as several WM properties appear to be affected. Moreover, the metrics most often used in other studies (i.e., FA and MD) were not identified as strong contributors in our study, highlighting the importance of using more physiologically-specific metrics from advanced imaging models as well as myelin-sensitive techniques. Further research combining advanced imaging metrics through multivariate approaches is needed to comprehensively evaluate the impact of CAD on brain WM microstructural health.

### Links with cognition

WM abnormalities have been implicated as a key mechanism underlying cognitive impairments associated with aging and vascular diseases (Filley & Fields, 2016; Vernooij et al., 2009; Chen et al., 2018; Santiago et al., 2015). Although executive function and processing speed did not significantly differ between groups in this study, we observed positive associations between processing speed and R1 in the left MCA territory as well as across the whole WM (Figure 3), suggesting that myelination differences may contribute to subtle individual variations in cognitive performance. This finding aligns with previous studies reporting associations between processing speed (i.e., reaction time) and WM microstructure in cognitively unimpaired older adults (Kerchner et al., 2012). Santiago and colleagues (2015) similarly reported that, in cognitively unimpaired patients with CAD, FA in the left parahippocampal cingulum and inferior fronto-occipital fasciculus was positively associated with a composite score combining executive function and processing speed. However, their findings were limited to individuals with CAD, as the study lacked a control group. In contrast, our study found that processing speed was the only cognitive domain that correlated with WM microstructure, and these associations were observed across all participants, regardless of CAD status. This suggests that the mechanism which underlies the link between R1 and processing speed is not specific to the disease, and can contribute to lower cognitive performance in healthy individuals as well.

The MCA territory is particularly vulnerable to ischemic stroke and CAD, likely due to its direct connection to the internal carotid artery, a common site of atherosclerosis (Vigen et al., 2020; Mathur et al., 1963). Consistent with this, we observed significantly higher D2 values in the bilateral MCA territory of individuals with CAD, indicating greater WM microstructural alterations in this region. Additionally, we found a positive association between processing speed and R1 in the left MCA. Although no significant group differences in processing speed were observed, the elevated D2 values in the MCA of participants with CAD, combined with the relationship between myelin content and processing speed, suggest that microstructural damage in this region may contribute to cognitive impairments in CAD. These findings reinforce the notion that WM alterations play a critical role in the early cognitive consequences of vascular dysfunction. Future research should further explore how WM microstructural integrity in watershed areas between cerebral arterial territories influences cognition.

### Limitations & Future directions

A strength of our study is the use of advanced models of WM microstructure that yield several measures of WH health, including measures of axonal density, neurite dispersion and myelin content. These measures were integrated using a multivariate approach developed by our group (Tremblay et al., 2024). The interpretation of this score as a measure of the extent of WM abnormality was facilitated by the presence of a well-defined HC group. Incorporating several WM metrics, reflecting various aspects of WM microstructure, offered a more comprehensive understanding of the underlying pathological mechanisms underlying WM alterations. This multivariate approach provided valuable insights by allowing the identification of WM metrics that are most affected by CAD. Additionally, we were the first to investigate WM microstructural health in cerebral arterial territories using a 3D MR-based atlas (Liu et al., 2023).

However, our study also suffered from some limitations. The small sample size gave us limited statistical power, which may have impaired our ability to detect associations between cognition and WM alterations. The sample size also did not allow the investigation of more complex interactions between variables, including the effects of sex. Future research in larger samples could evaluate WM microstructure and cognition across participants with a broader spectrum of cardiovascular health, for instance, by recruiting participants with cardiovascular risk factors. This would provide a better understanding of the mechanisms through which cardiovascular health impacts the brain from the earliest stage.

## Conclusion

This study employed a novel multivariate approach to assess WM health in both HCs and individuals with CAD. Our findings revealed widespread WM abnormalities in patients with CAD, affecting both global WM and specific arterial territories. Additionally, myelination in the WM of the left MCA territory and the whole WM was associated with processing speed across all participants. These results suggest that the higher WM abnormalities observed in CAD may contribute to a heightened risk of cognitive impairment. Future research should explore whether interventions such as cardiac rehabilitation—incorporating exercise, dietary modifications, and cognitive training—can preserve or restore WM integrity in individuals at risk for or diagnosed with CAD. Advancing research in this area is crucial, as targeted interventions may help prevent the progression from subclinical vascular disease to vascular cognitive impairment and dementia (Inoue et al., 2023; Poirier et al., 2024; Middleton et al., 2008; Moorhouse & Rockwood, 2008).

## Acknowledgments

We would like to thank everyone who contributed to this project: Paule Samson, Thomas Vincent, Julie Lalongé, Hakima Benhalima, Milla Shakleva, Victoria D’Amours, Agathe Godet, Stephanie Beram, Roni Zaks, Robert Hovey, Alexandre Bailey, Catherina Medeiros, and Zineb Rouabah. Thank you also to the laboratories of Dr Louis Bherer and Dr Mathieu Gayda. We would also like to acknowledge the McConnell Brain Imaging Centre of the Montréal Neurological Institute, which provided a custom pulse sequence for the MTI protocol. Lastly, we would like to acknowledge our research participants, without whom none of this would have been possible.

