## Supplemental Figure 1 & Table 1 for "Multivariate white matter microstructure alterations in older adults with coronary artery disease"

Supplemental Material

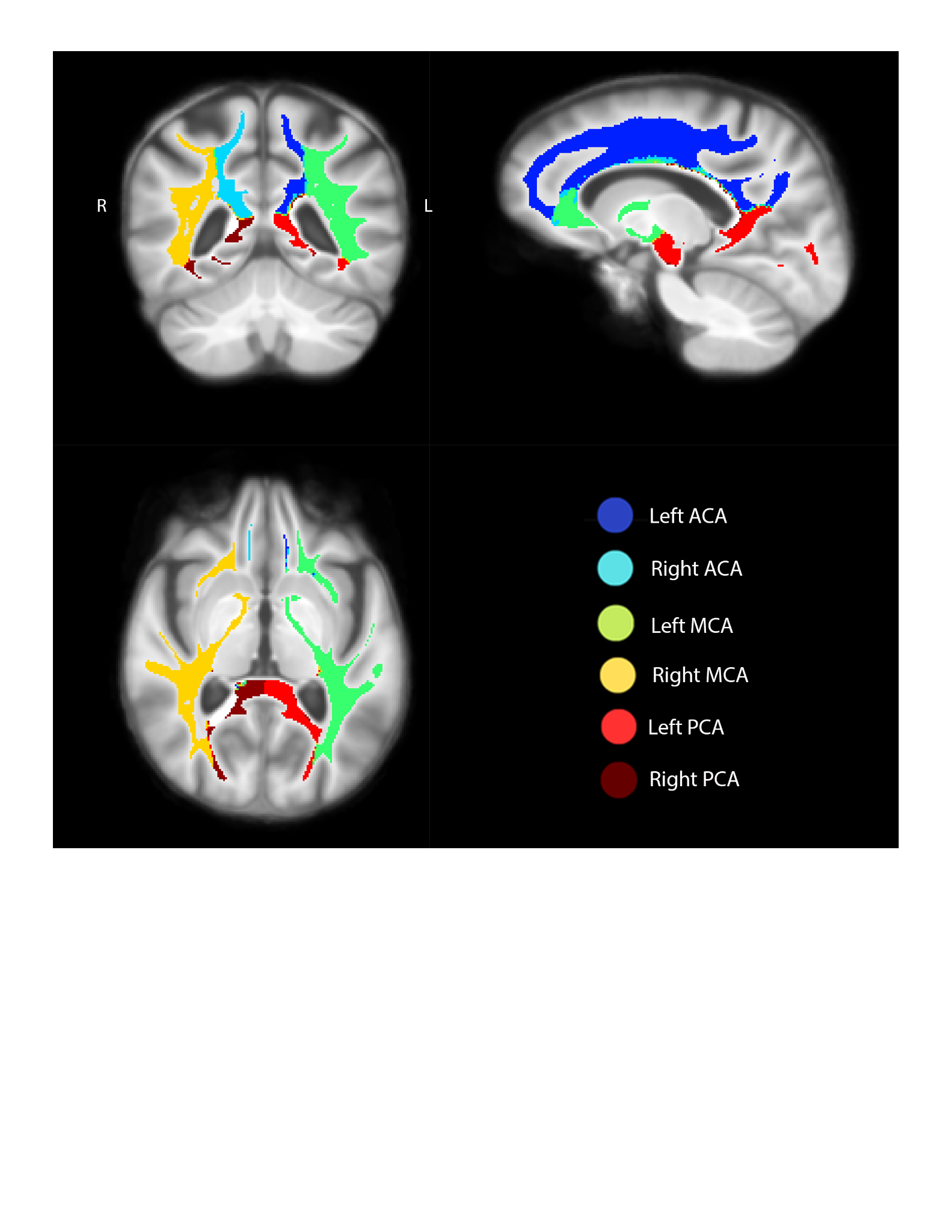

**Figure S1**. Arterial territories within the WM mask, overlaid on the group average MPRAGE T1w image.

**Table S1.** WM metrics contributions in each region of interest (only those that contributed >10% to D2 are shown**)**.

| **Regions of Interest** | **Percentage contribution to D2** |
| --- | --- |
| Left ACA | R1 (13.4%), ICVF (12.3%), FA (10.7%) |
| Right ACA | R1 (18.0%), MTsat (10.3%) |
| Left MCA | R1 (14.1%), RD (12.0%), MTsat (10.7%) |
| Right MCA | R1 (15.0%), MTsat (14.5%), RD (10.7%) |
| Left PCA | R1 (21.7%), MTsat (10.5%) |
| Right PCA | R1 (23.0%), FA (10.4%) |

*ACA = Anterior Cerebral Artery; MCA = Middle Cerebral Artery; PCA =Posterior Cerebral Artery; ICVF = Intracellular Volume Fraction; RD = Radial Diffusivity; FA = Fractional Anisotropy*
